# Video diffusion for early cellular apoptosis forecasting

**DOI:** 10.1101/2023.11.16.567461

**Authors:** Akash Awasthi, Jaer Nizam, Samira Zare, Safwan Ahmad, Melisa J. Montalvo, Navin Varadarajan, Badri Roysam, Hien Nguyen

**Affiliations:** University of Houston

**Keywords:** Video Diffusion, Apoptosis, Cellular Activity Forecasting

## Abstract

Reliable and early prediction of cell death (apoptosis) is critically important in various areas of biology, particularly in characterizing the effectiveness of cell-based infusion products utilized for cancer immunotherapy. While deep Convolutional Neural Networks (CNNs) are often used in state-of-the-art approaches for apoptosis classification, they typically focus solely on individual cells and ignore cell-cell interaction. To address this limitation, we propose a novel generative approach based on a video diffusion model, which predicts future cellular behaviors for early detection of apoptosis events, even before molecular markers like Annexin-V or visual indications like membrane blebbing become apparent. Our approach accounts for the interactions of multiple target cells and their spatial and temporal relationships at each time frame. We condition our generative model on two starting frames and utilize an auto-regressive framework to predict the subsequent five frames. Our model achieves a 0.88 F1 score on the cell death event classification and 2.11 mean absolute death-time error, significantly outperforming state-of-the-art methods.

## 1. INTRODUCTION

### 1.1. Cell Death and Diffusion Models

In recent years, immunotherapy has been a transformative breakthrough in cancer treatment. One such treatment includes T cell therapy, which showed promise in killing cancer cells. Researchers emphasized the significant progress made in tumor eradication using engineered T cells such as CAR-T, which induce apoptosis to eliminate tumor cells [10]. In [22], it can be seen how anticancer compounds also induced apoptosis to eliminate cancer cells. Evidently, we can see that apoptosis will play a significant role in future innovations and advancements in cancer therapy. Apoptosis is the normal and controlled cell death event that occurs in multicellular organisms as part of an organism’s growth or development[2]. The process of apoptosis includes membrane blebbing, cell shrinking, deoxyribonucleic acid (DNA) degradation, and in some cases the formation of apoptotic bodies that depend on the size of target cells[3]. Recently, there has been a high demand for automated identification or categorization of apoptosis due to the emergence of high throughput screening assays [17, 9]. Additionally, advancements in immunotherapy for cancer treatment have resulted in the development of dynamic cell-based assays such as TIMING (time-lapse microscopy in nanowell grids) [13]. These assays have enabled the monitoring of immune cell-cancer cell interactions in numerous sub-nanoliter wells, offering valuable insights into the biology of immune cells and their behavior towards tumor cells in various pre-clinical scenarios [6, 14].

Apoptosis is a complex process that unfolds over multiple image frames. There is a compelling need to detect and quantify all aspects of this process to evaluate the effectiveness of experimental therapeutics. Since apoptosis occurs in a subset of nanowells containing target and effector cells, it is valuable to identify the nanowells in which it is most likely to occur to focus the measurement efforts.

In this method, we propose a novel apoptosis prediction framework, called video Diffusion for Early Cellular Apoptosis Forecasting (DECAF). Diffusion models are generative models that give insight into how information is propagated through a neural network. These models attempt to distort the training data by adding noise to it. Subsequently, it tries to reverse the effects of the noise to recover the original data. From the noise levels that it estimates, the model can learn the probability distribution of the data set, allowing the model to generate its own realistic samples of the data.

Our paper makes the following contributions:

1. We propose to use a video diffusion model to predict apoptosis early by generating the phase-contrast and florescent channel simultaneously.
2. The model generates images of cells and nanowells. By analyzing these frames our model categorizes cell death events and captures cell interactions to visualize how cell communication contributes to apoptosis.

### 1.2. Video Diffusion Models

Diffusion models have been previously used for image generation. Nichol et al. [16] uses the clip-guided generation for modeling the conditional distribution of data. Some of the score-based diffusion models are used for video generation in [20]. Ho et al. [5] proposed the video diffusion model which uses the gradient method to approximate the conditional distributions. Other models such as RVD (Residual Video Diffusion) applied by Yang et al. [24] were used on video prediction. RVD models the residual of the predicted and true video frames. This model was able to model conditional distribution but only considers past frames as conditions.

## 2. RELATED WORKS

Since CNN models lacked the ability to generate videos and pictures of the whole cell environment; video diffusion models were applied to simulate the different cell interactions that exist during a cell death event. Using standard Gaussian diffusion models, Ho et al. [5] first introduced the application of diffusion models in video modeling. Voleti et al. [21] displayed the use of video diffusion models through a framework they constructed called ‘Masked Conditional Video Diffusion’(MCVD). This framework had state-of-the-art performance in video synthesis tasks such as future/past prediction, unconditional generation, and interpolation. Dockhorn et al. [1] introduces a second-order ODE solver called ‘GENIE’. This solver truncates Taylor methods to solve the generative ODE of denoising diffusion models with higher accuracy than previous methods. The research by Song et al. [19] presents another method of denoising for diffusion models. They introduce ‘denoising diffusion implicit models’(DDIMs) as a way to close the efficiency gap between generative adversarial networks (GANs) and denoising diffusion probabilistic models (DDPMs), which require many iterations to produce high-quality samples. Ramesh et al. [18] combines the well-known Contrastive Language-Image Pre-Training (CLIP), by OpenAI, with generative diffusion models to train a model that can demonstrate the ability to produce semantically similar output images and interpolate between images.

## 3. METHODOLOGY

### 3.1. Conditional Diffusion for Video

Given ***x***_0_ ∼ *q*(***x***), the forward process corrupts ***x***_0_ with a small amount of Gaussian noise at each time steps *t* ∈ [0, *T*] that satisfies Markovian transition:

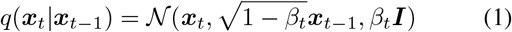

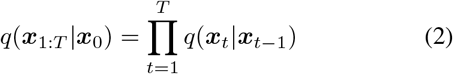

The initial image ***x***_0_ is gradually degraded as the step becomes larger and eventually, ***x***_*T*_ is equivalent to an isotropic Gaussian distribution. An elegant property of this process is that from ***x***_0_, ***x***_*t*_ any arbitrary *t* can be sampled using the reparameterization technique [8] as follows:

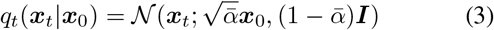

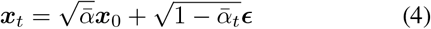

where 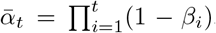, and ***ϵ*** ∼ *𝒩*(**0, *I***). To reverse the above process and generate new samples, we need to approximate *q*(***x***_*t*−1_|***x***_*t*_) by learning *p*_*θ*_, which is tractable when conditioned on ***x***_0_:

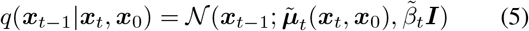

where

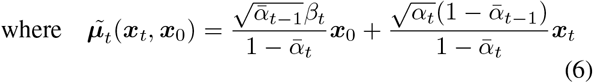

and

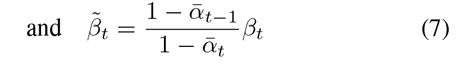

Thanks to this nice property, we can 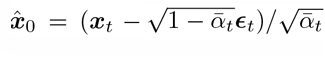. Since ***x***_*t*_ is available from the forward process, we can re-parameterize the Gaussian noise term to predict ***ϵ***_*t*_ by *p*_*θ*_. Thus, the loss term would be:

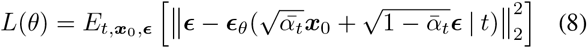

### 3.2. Video Prediction Using Conditional Diffusion

We model the conditional distribution of the video frames by incorporating the past frames. The diffusion model is conditioned on past frames to predict future frames.

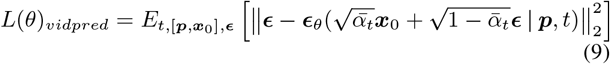

After training the diffusion model, ***ϵ***_*θ*_, shown in Equation 9, is modeled using variants of neural networks shown in Fig. 2. In the first window of *p* + *k* frames, we predict the *k* frames conditioned on the past *p* frames, and then we shift the window to the *p* + *k* frames and repeat the same process for the whole video.

### 3.3. Architecture

Our model is based on the video prediction architecture proposed by Voleti et al. [21]. To denoise the input frames, we used a U-Net backbone with several modifications. The architecture includes multi-head self-attention, 2D convolution, and adaptive group normalization [23]. Additionally, we used transformer-style encoding to process the position encoding of the noise level.

Our model utilizes past frames as conditioned frames, which are concatenated along the channel dimension. In the forward diffusion process, current noisy frames are created and used as input for the denoising network at time step *t*. The concatenated conditional frames are passed through a network that employs Spatially-Adaptive (DE)normalization (SPADE), which accounts for both time and motion. This approach is preferable to using Flownet, which would increase computational complexity.

For each embedding vector, **e**(*t*), we apply the following formula:

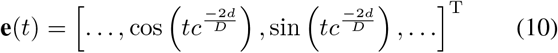

Here *d* = 1, …, *D/*2, *D* is the number of dimensions of the embedding, and *c* = 10000. The resulting embedding vector is passed through a fully connected layer with an activation function, followed by another fully connected layer.

### 3.4. Data Sets

#### Cell Culture and Imaging

The cell images used in this study were obtained from four TIMING data sets. We used the fluorescent membrane stains, PKH67 green (Sigma-Aldrich) and PKH26 red (Sigma-Aldrich), to label CAR T-cells and NALM6 cells, respectively. Both cells were then loaded into nanowells at a concentration of 1 million cells/mL.The cells were incubated at a constant temperature of 37 °C and 5% CO2 for the duration of imaging. The cells were imaged using phase contrast, Alexa Fluor 488, Texas-Red, and CY5 filters using a Carl Zeiss Observer Z1 fitted with a Hamamatsu sCMOS camera and a 20 × 0.8 NA objective lens for six hours at 5-minute intervals. Fig. 1 shows the structure of a nanowell array and frames captured from a nanowell demonstrating an apoptotic cell. These results provide valuable insights into the classification of cell apoptosis and may aid in future studies.

**Fig. 1:**
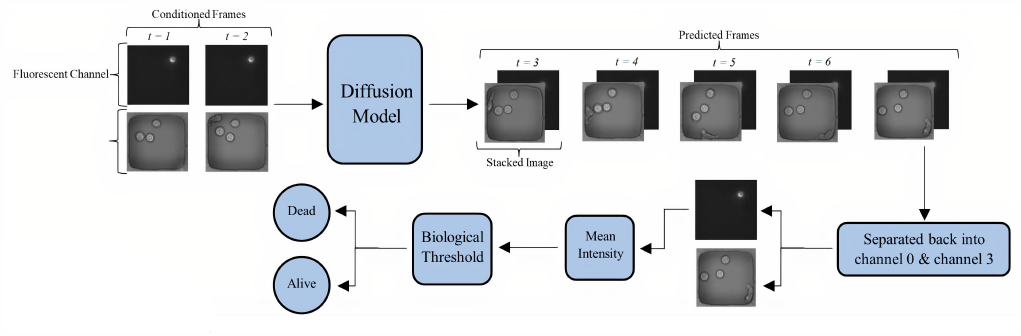
Overview of the proposed diffusion model for cell video generation. Generated videos are used to classify or forecast apoptotic events.

**Fig. 2:**
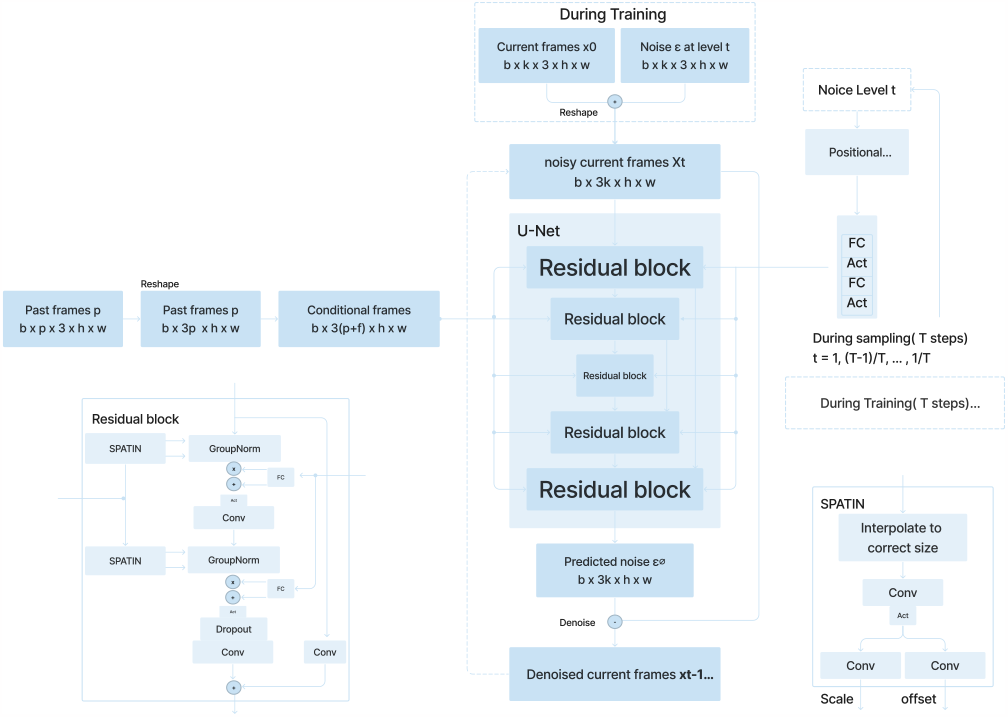
A U-Net model that denoise current frames by predicting the noise in the frames based on past/future frames and noise level.

#### TIMING Pipeline and the Ground Truth

The TIMING pipeline is a collection of algorithms designed for nanowell detection, cell segmentation, and cell tracking. With this tool, we were able to analyze the entire nanowell array at each time point, rather than examining each cell individually. The TIMING pipeline produces four channels for each time point, including phase contrast, effector cell fluorescence, target cell fluorescence, and death marker channels.

To classify the data for apoptosis, we used a stacked image of the phase contrast channel and fluorescent channels, as shown in Fig. 1. We employed a binary labeling system, labeling the nanowell array with ‘0’ indicating that the cell was alive, or with a ‘1’ indicating that it was dead. We used a simple threshold method to label the data. The mean intensity of each fluorescent channel was calculated, and then this value was compared to a biological threshold set by experts in the field. If the mean intensity was lower than the threshold, the nanowell was labeled as 1; if the mean intensity was greater than the threshold, the nanowell was labeled as 0. We assumed that once a nanowell was identified as dead, it remained dead for all subsequent frames.

## 4. EXPERIMENTAL RESULTS

We have used the diffusion model architecture for the video prediction task. We use the 2 past frames as conditioned frames and predict 5 frames at a time and then shift to the next window. For sampling, we use the DDPM sampling [4] with the 100 sampling steps but the model was trained with 100 sampling steps. We have stacked the florescent channel, phase contrast channel, and image of zeros to form a three-dimensional array. This three-dimensional array is used as input to the diffusion model as shown in the figure. The model predicts the three-dimensional image. The fluorescent channel and phase contrast channel are separated from this three-dimensional input.

### 4.1. Apoptosis Classification

The proposed diffusion model is used for generating the cell dead and alive events. We compute its performance on cell death classification. We compared its performance with the GAN-based method [11], and the convolution-LSTM–based method. All networks are trained using images of size 61 × 61 pixels. Training is done using the ADAM optimizer [7] with an initial learning rate of 10–4 and mini-batches of size 2. All the model weights are initialized using Xavier initialization. The ROC-AUC metric and accuracy metric are compared with the two methods mentioned above. Table 1 shows the same.

**Table 1:** AUC Score Comparison with three future frames.

| Model (Frame number) | ROC-AUC | Accuracy | MADTE |
| --- | --- | --- | --- |
| DECAF (1) | 0.88 | 80% | 2.11 |
| DECAF (2) | 0.83 | 72% | 2.35 |
| DECAF (3) | 0.84 | 75% | 2.31 |
| Future GAN [11] (1) | 0.61 | 73% | 3.44 |
| Future GAN [11] (2) | 0.57 | 70% | 3.71 |
| Future GAN [11] (3) | 0.58 | 71% | 3.62 |
| ConvLSTM [12] (1) | 0.65 | 72% | 4.17 |
| ConvLSTM [12] (2) | 0.63 | 70% | 7.3 |
| ConvLSTM [12] (3) | 0.60 | 69% | 8.1 |

The confusion metrics in fig 3 and ROC-AUC is calculated for the diffusion model for cell death classification. It also shows the learning curve for diffusion model which converges well.Our model is able to classify cell death events with and ROC-AUC of 0.82. This level of performance is promising, considering that we do not use any ground truth labels during training. Our model significantly outperforms state-of-the-art methods based on GAN [11] (0.61 ROC-AUC) and ConvLSTM [12] (0.65 ROC-AUC).

**Fig. 3:**
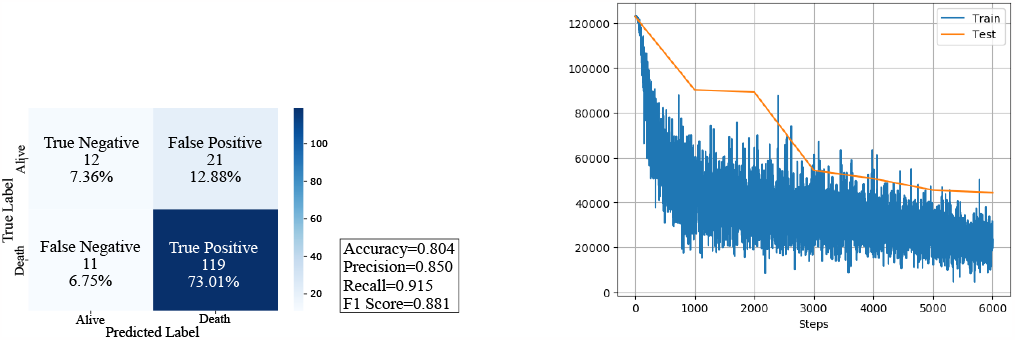
Confusion matrix/Learning curve for Diffusion model

### 4.2. Early Apoptosis Forecasting

We have used two conditional frames and predicted five frames (25 minutes ahead) at a time using the sliding window approach. Fig 4 shows the real and predicted cell images. The first row shows the real images for video-1, and the second row shows the early prediction of cell death events. The model predicts the cell death event and cell dynamics very well. The second row shows video-2.

**Fig. 4:**
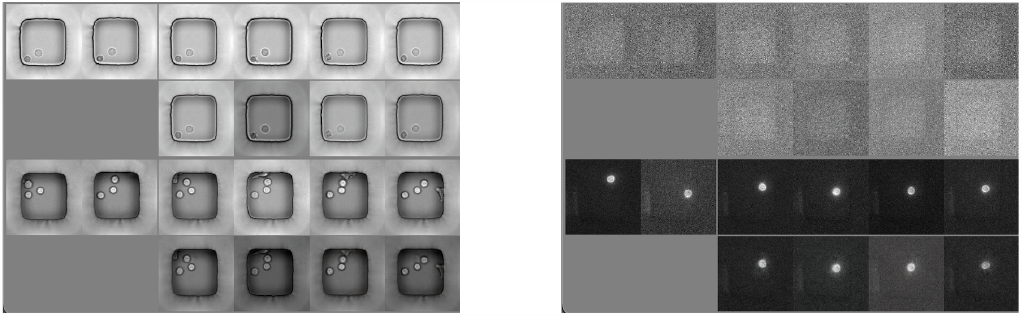
Early detection of cell death event in phase contrast channel (left) and fluorescent channel (right) for early detection of cell death event. The row 1 and row 3 show the true frames, while row 2 and row 4 show the predicted frames.

Fig. 4 also shows the prediction of the third channel. We separate the phase contrast and fluorescent channel from the predicted images. As can be seen in the figure, the model predicts the death marker for the next 8 channels which shows that this video represents the cell death event.

### 4.3. Cell death time detection

In order to calculate the accuracy of the model in detecting the death time which informs how early the cell death is predicted. This is measured by the mean absolute death-time error (MADTE) [15] computed as:

The value we get for our model is 2.11 which is less compared to other methods as reported in the table 1

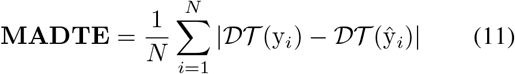

where *𝒟 𝒯*(.) :R^*L*^ →R is the function that returns the death time, i.e. index of the first frame labeled ‘l’.

## CONCLUSIONS

Our work shows that a generative model can be used to perform novel and surprising types of biological image analysis. Our experiments show that a complex phenomenon like apoptosis can be predicted with good accuracy even before the images recording apoptosis are collected. This is in contrast to traditional image analysis methods for detecting cellular events by analyzing collected images that become available after the events have occurred. Our experiments suggest that generative modeling is able to capture complex aspects of biological phenomena, even at a multi-cellular scale. In future work, we expect to generate interpretations of the generative models that can provide a direct path from image collection to biological model discovery.

## REFERENCES

[1] Tim Dockhorn, Arash Vahdat, and Karsten Kreis. Genie: Higher-order denoising diffusion solvers. arXiv preprint arXiv:2210.05475, 2022.

[2] Susan Elmore. Apoptosis: a review of programmed cell death. Toxicologic pathology, 35(4):495–516, 2007.

[3] Yaron Fuchs and Hermann Steller. Programmed cell death in animal development and disease. Cell, 147(4):742–758, 2011.

[4] Jonathan Ho, Ajay Jain, and Pieter Abbeel. Denoising diffusion probabilistic models. Advances in Neural Information Processing Systems, 33:6840–6851, 2020.

[5] Jonathan Ho, Tim Salimans, Alexey Gritsenko, William Chan, Mohammad Norouzi, and David J Fleet. Video diffusion models. arXiv preprint arXiv:2204.03458, 2022.

[6] Carl H June and Michel Sadelain. Chimeric antigen receptor therapy. New England Journal of Medicine, 379(1):64–73, 2018.

[7] Diederik P Kingma and Jimmy Ba. Adam: A method for stochastic optimization. arXiv preprint arXiv:1412.6980, 2014.

[8] Durk P Kingma, Tim Salimans, and Max Welling. Variational dropout and the local reparameterization trick. Advances in neural information processing systems, 28, 2015.

[9] Jonas Kühn, Etienne Shaffer, Julien Mena, Billy Breton, Jérôme Parent, Benjamin Rappaz, Marc Chambon, Yves Emery, Pierre Magistretti, Christian Depeursinge, et al. Label-free cytotoxicity screening assay by digital holographic microscopy. Assay and drug development technologies, 11(2):101–107, 2013.

[10] D. Li, X. Li, WL. Zhou, and et al. Genetically engineered t cells for cancer immunotherapy. Signal Transduction and Targeted Therapy, 4(35):1–10, 2019.

[11] Wen Liu, Weixin Luo, Dongze Lian, and Shenghua Gao. Future frame prediction for anomaly detection–a new baseline. In Proceedings of the IEEE conference on computer vision and pattern recognition, pages 6536–6545, 2018.

[12] Weixin Luo, Wen Liu, and Shenghua Gao. Remembering history with convolutional lstm for anomaly detection. In 2017 IEEE International Conference on Multimedia and Expo (ICME), pages 439–444. IEEE, 2017.

[13] Amine Merouane, Nicolas Rey-Villamizar, Yanbin Lu, Ivan Liadi, Gabrielle Romain, Jennifer Lu, Harjeet Singh, Laurence JN Cooper, Navin Varadarajan, and Badrinath Roysam. Automated profiling of individual cell–cell interactions from high-throughput time-lapse imaging microscopy in nanowell grids (timing). Bioinformatics, 31(19):3189–3197, 2015.

[14] Androulla N Miliotou and Lefkothea C Papadopoulou. Car t-cell therapy: a new era in cancer immunotherapy. Current pharmaceutical biotechnology, 19(1):5–18, 2018.

[15] Aryan Mobiny, Hengyang Lu, Hien V Nguyen, Badrinath Roysam, and Navin Varadarajan. Automated classification of apoptosis in phase contrast microscopy using capsule network. IEEE transactions on medical imaging, 39(1):1–10, 2019.

[16] Alex Nichol, Prafulla Dhariwal, Aditya Ramesh, Pranav Shyam, Pamela Mishkin, Bob McGrew, Ilya Sutskever, and Mark Chen. Glide: Towards photorealistic image generation and editing with text-guided diffusion models. arXiv preprint arXiv:2112.10741, 2021.

[17] Anna-Liisa Nieminen, Gregory J Gores, John M Bond, Roberto Imberti, Brian Herman, and John J Lemasters. A novel cytotoxicity screening assay using a multiwell fluorescence scanner. Toxicology and applied pharmacology, 115(2):147–155, 1992.

[18] Aditya Ramesh, Prafulla Dhariwal, Alex Nichol, Casey Chu, and Mark Chen. Hierarchical text-conditional image generation with clip latents. arXiv preprint arXiv:2204.06125, 2022.

[19] Jiaming Song, Chenlin Meng, and Stefano Ermon. Denoising diffusion implicit models. arXiv preprint arXiv:2010.02502, 2020.

[20] Ruben Villegas, Jimei Yang, Seunghoon Hong, Xunyu Lin, and Honglak Lee. Decomposing motion and content for natural video sequence prediction. arXiv preprint arXiv:1706.08033, 2017.

[21] Vikram Voleti, Alexia Jolicoeur-Martineau, and Christopher Pal. Masked conditional video diffusion for prediction, generation, and interpolation. arXiv preprint arXiv:2205.09853, 2022.

[22] Jack H Wong, Stephen CW Sze, Tzi Bun Ng, Randy Chi Fai Cheung, Chit Tam, Kalin Yanbo Zhang, Xiuli Dan, Yau Sang Chan, William Chi Shing Cho, Charlene Cheuk Wing Ng, et al. Apoptosis and anti-cancer drug discovery: the power of medicinal fungi and plants. Current Medicinal Chemistry, 25(40):5613–5630, 2018.

[23] Y Wu and K He. Group normalization. arxiv. arXiv preprint arXiv:1803.08494, 2018.

[24] Ling Yang, Zhilong Zhang, Yang Song, Shenda Hong, Runsheng Xu, Yue Zhao, Yingxia Shao, Wentao Zhang, Bin Cui, and Ming-Hsuan Yang. Diffusion models: A comprehensive survey of methods and applications. arXiv preprint arXiv:2209.00796, 2022.

